# Predicting the toxicity of chemical compounds via Hyperdimensional Computing

**DOI:** 10.1101/2025.09.12.675894

**Authors:** Fabio Cumbo, Kabir Dhillon, Jayadev Joshi, Bryan Raubenolt, Davide Chicco, Sercan Aygun, Daniel Blankenberg

**Author notes:** To whom correspondence should be addressed: Daniel Blankenberg^1,6^, Center for Computational Life Sciences, Cleveland Clinic Research, Cleveland Clinic, 9500 Euclid Avenue, NA2, Cleveland, OH 44195, USA.

## Abstract

Accurately and efficiently assessing the potential toxicity of chemical compounds is critical given their wide application across pharmaceutical, industrial, and environmental domains. Traditional toxicological evaluations, which predominantly rely on intensive *in vitro* and *in vivo* assays, are frequently slow and expensive processes. Here, we introduce a novel application of Hyperdimensional Computing (HDC), an emerging computational paradigm inspired by the way the human brain works in encoding information, for the efficient classification of chemical compounds as either toxic or non-toxic. Our methodology employs Simplified Molecular Input Line Entry System (SMILES) representations of compounds, drawing data from the comprehensive Tox21 dataset. We delineate a pipeline wherein these chemical structures are encoded into high-dimensional binary vectors, which subsequently serve as the foundation for training and classification within the HDC framework. This approach leverages HDC’s inherent advantages, including its resilience to noise, parallel processing capabilities, and efficacy in identifying intricate patterns. This work demonstrates the viability of HDC as a promising alternative for large-scale toxicity prediction, offering a computationally efficient and scalable solution. This research significantly contributes to the field of cheminformatics by validating HDC’s potential in chemical property prediction, thereby facilitating accelerated identification of hazardous substances and mitigating the reliance on intensive laboratory experimentations.

## 1. INTRODUCTION

The study and discovery of new chemical compounds across modern society is unprecedented [1]. From life-saving pharmaceuticals and crop-enhancing agrochemicals to industrial solvents and everyday consumer products, chemical compounds are integral to countless facets of daily life [2,3]. However, this ubiquity carries an inherent risk: the potential for adverse effects on human health and the environment. The huge volume of new and existing compounds entering the market vastly outspaces our capacity to rigorously evaluate their safety. Consequently, accurately and efficiently assessing the potential toxicity of these substances has become one of the most pressing challenges in public health, environmental science, and regulatory oversight [4].

Traditionally, toxicological assessment has depended on a paradigm of *in vitro* and *in vivo* assays [5]. While these methods have long been the gold standard, they are affected by significant limitations. They are notoriously slow, often requiring months or even years to complete for a single compound. Furthermore, they are exceptionally expensive and resource-intensive, demanding significant investment in laboratory infrastructure, materials, and specialized personnel [6]. Beyond the practical constraints, the reliance on animal testing raises substantial ethical concerns, prompting a global push towards the development of alternative, non-animal-based testing strategies [7]. These bottlenecks create a critical gap in safety data, leaving regulators, manufacturers, and the public with an incomplete understanding of the potential hazards associated with the vast number of chemicals currently in circulation.

To bridge this gap, the field of computational toxicology has emerged, seeking to leverage computational power to predict the biological activity of chemicals based on their molecular structure. These *in silico* methods offer a powerful alternative, promising rapid, cost-effective, and large-scale screening of chemical libraries [8–10]. Prominent among these approaches are Quantitative Structure-Activity Relationship (QSAR) models, which establish mathematical relationships between a compound’s structural or physiochemical properties (known as molecular descriptors) and its toxicological endpoint [11,12]. In recent years, various machine learning algorithms have been successfully applied to this problem, often demonstrating high predictive accuracy [13,14]. However, many of these conventional methods depend on a crucial and often complex pre-processing step: the calculation of handcrafted molecular descriptors. This process of feature engineering could eventually affect the model’s performance because of being limited by the scope of the pre-selected features, a process that can inadvertently discard subtle but important structural information encoded within the molecule [15].

This paper explores a fundamentally different computational paradigm called Hyperdimensional Computing (HDC) [16–18], also known as Vector-Symbolic Architectures (VSA), for toxicity prediction. It is an emerging paradigm inspired by the distributed, robust, and efficient nature of information processing in the human brain with growing applications in biomedical sciences [19]. Instead of operating on precise, low-dimensional numbers, HDC represents information using very high-dimensional vectors, often referred to as hypervectors, typically with thousands of elements. In this framework, information is distributed across the entire vector, making the representation inherently resilient to noise and errors [20]. HDC employs a simple yet powerful algebra of vector operations to construct simple representations of complex structures by performing cognitive tasks like learning, classification, and reasoning [21–23].

These principles make it particularly suitable to the challenges of cheminformatics. Chemical structures, as represented by notations like the Simplified Molecular-Input Line-Entry System (SMILES) [24], are inherently sequential and compositional. The HDC framework provides a natural mechanism for encoding such *symbolic data* into high-dimensional space without relying on pre-computed descriptors as in the study by Ma *et al*. [25] from which this study takes inspiration. Our work builds upon this foundation by providing a more focused and methodologically deep investigation. While the work of Ma *et al*. also explored atom-wise and k-mer tokenization, our study extends this analysis by introducing a chemically-informed, fragment-base approach, enabling a broader comparison across different levels of structural granularity. Furthermore, to the best of our knowledge, this is the first study to apply the HDC paradigm specifically to the problem of toxicity prediction. We do so by leveraging the comprehensive and challenging multi-target Tox21 dataset [26], implementing a robust validation protocol to handle its severe class imbalance. Because HDC is well-suited to handling raw input values with a single-pass learning strategy, extensive pre-processing or feature engineering is not always required. By directly processing SMILES strings, an HDC model can learn to represent and associate complex structural motifs and their arrangements, capturing the patterns that determine a compound’s biological activity. This approach leverages HDC’s inherent strengths in pattern recognition, its scalability, and its ability to operate efficiently on raw, unprocessed data.

In this study, we introduce and validate a novel HDC-based pipeline for classifying chemical compounds as toxic or non-toxic. The remainder of this manuscript is organized to systematically present our investigation. The next section details the full computational methodology, beginning with the acquisition and preparation of the dataset. We then elaborate on the crucial step of transforming chemical structures into machine-readable formats by comparing different SMILES tokenization strategies. Subsequently, we outline the design of our HDC model, from the fundamental principles of VSAs to the specific encoding schemes and classification components used. We then present the empirical results, evaluating the performances of our approach in predicting toxicity across different biological targets. Finally, we provide a discussion of these results and their broader implications, followed by a conclusion that summarizes our key findings and suggests avenues for future work.

## 2. MATERIALS AND METHODS

Here, we delineate the experimental design and computational procedures adopted to investigate the efficacy of HDC for predicting the toxicity of chemical compounds, starting with data acquisition, different tokenization strategies, model construction, to conclude with a test and evaluation protocol.

### 2.1. Dataset acquisition and preprocessing

This study is based on developing novel predictive toxicity models by leveraging the Tox21 dataset, a publicly available collection of chemical compounds and their associated toxicity labels [26]. This dataset originates from the Tox21 program, a collaborative effort by federal agencies to identify compounds that disrupt human biological pathways. The dataset structure is characterized by two primary components for each chemical entry: a SMILES string representing the compound structure and 12 binary columns indicating its interaction with specific biological receptors. A value of 1 under a receptor column for a specific compound indicates an interaction, 0 otherwise. In this context, a compound is considered toxic if it shows an interaction with any of the 12 receptors.

A total of 7,832 compounds are present in this dataset. However, it is important to note that not all compounds have a defined outcome for all 12 receptors. The dataset contains missing values, which signify that the original experiment for a specific compound-receptor pair was inconclusive (e.g., due to noisy data or an ambiguous response). Consequently, the total number of compounds (toxic + non-toxic) analyzed for each individual receptor is typically less than the total number of compounds reported in the Tox21 dataset.

The 12 receptors included in the Tox21 dataset, along with their brief descriptions, are summarized in Table 1.

**Table 1:** List of the 12 biological receptors used in the Tox21 dataset to differentiate toxic and non-toxic chemical compounds.

| Receptor | Toxic compounds | Non-toxic compounds | Description |
| --- | --- | --- | --- |
| NR-AR | 309 | 6,956 | <u>Nuclear Receptor – Androgen Receptor</u>: involved in male sexual development and function, often targeted by endocrine disruptors. |
| NR-AR-LBD | 237 | 6,521 | <u>Nuclear Receptor – Androgen Receptor Ligand Binding Domain</u>: specifically targets the ligand-binding domain of the androgen receptor, a common site for chemical interactions. |
| NR-AhR | 768 | 5,781 | <u>Nuclear Receptor – Aryl Hydrocarbon Receptor</u>: a ligand-activated transcription factor involved in the regulation of genes in response to environmental pollutants. |
| NR-Aromatase | 300 | 5,521 | <u>Nuclear Receptor – Aromatase</u>: an enzyme responsible for a key step in the biosynthesis of estrogens, inhibition of which can have endocrine effects. |
| NR-ER | 793 | 5,400 | <u>Nuclear Receptor – Estrogen Receptor alpha</u>: that is activated by the hormone estrogen, playing a role in female reproductive function and other processes. |
| NR-ER-LBD | 350 | 6,605 | <u>Nuclear Receptor – Estrogen Receptor Ligand Binding Domain</u>: focuses on the ligand-binding domain of the estrogen receptor, a critical site for endocrine disruption. |
| NR-PPAR-gamma | 186 | 6,264 | <u>Nuclear Receptor – Peroxisome Proliferator-Activated Receptor gamma</u>: a nuclear receptor that plays a central role in adipogenesis, lipid metabolism, and glucose homeostasis. |
| SR-ARE | 942 | 4,890 | <u>Stress Response – Antioxidant Response Element</u>: a DNA regulatory element that controls the expression of a battery of genes involved in the cellular defense against oxidative stress. |
| SR-ATAD5 | 264 | 6,808 | <u>Stress Response – ATPase Family AAA Domain Containing 5</u>: involved in DNA repair pathways, and its disruption can lead to genomic instability. |
| SR-HSE | 372 | 6,095 | <u>Stress Response – Heat Shock Factor Response Element</u>: a DNA sequence that regulates the expression of heat shock proteins, crucial for cellular stress response. |
| SR-MMP | 918 | 4,892 | <u>Stress Response – Mitochondrial Membrane Potential</u>: assesses the integrity of mitochondrial function, as disruption of membrane potential can indicate cellular toxicity. |
| SR-p53 | 423 | 6,351 | <u>Stress Response – p53 Pathway</u>: a critical tumor suppressor protein that regulates cell cycle arrest, apoptosis, and DNA repair in response to cellular stress. |

### 2.2. SMILES Tokenization

Here, we introduce different techniques employed for transforming chemical structures, represented as SMILES strings into a tokenized format for subsequent hypervector encoding. SMILES is a line notation that allows a chemical structure to be represented by a short ASCII string. It provides a compact and unambiguous way to describe molecular graphs, encoding information about atoms, bonds, and molecular topology [24]. For each chemical compound in the Tox21 dataset, its unique SMILES string served as the primary input for our tokenization and encoding pipeline. Unlike traditional approaches that rely on pre-computed molecular descriptors [14], our method directly processes the SMILES strings, leveraging different tokenization granularities to explore their impact on HDC performance.

In order to capture diverse structural information from the SMILES strings, three distinct tokenization strategies were implemented and compared. Each strategy decomposes the SMILES string into a sequence of fundamental units, or tokens, which then form the basis for hypervector generation:

- <u>Atom-wise</u>: each individual atom and special SMILES character (e.g., =, #, (,), [,]) was treated as a distinct token. This method provides the most granular representation of the chemical structure, preserving information down to the level of individual atomic species and bonding types. For example, the SMILES string *CC*(= *O*)*O* would be tokenized into *C, C*, =, *O*,), *O*. This approach ensures that every atomic and structural detail explicitly represented in the SMILES string contributes to the token set;
- <u>K-mer-based</u>: this involves segmenting the SMILES strings into overlapping sequences of characters of a fixed length *k*. This method is analogous to n-gram approaches used in natural language processing, allowing the capture of local structural patterns within the chemical compounds. For instance, with *k* = 2, the SMILES *CC*(= *O*)*O* would yield tokens such as *CC, C*(, (=, = *O*,)*O*. This approach helps in identifying common motifs and their immediate chemical environment.
- <u>Fragment-based</u>: this tokenization method aims to capture chemically meaningful substructures or functional groups as tokens. This approach leverages predefined chemical fragments that represent common building blocks of molecules. For example, a carbonyl group *C*(= *O*) or a benzene ring *c*1*ccccc*1 could be recognized as single tokens. This strategy provides a higher level representation compared to atom-wise and k-mer tokenization, focusing on the presence of known chemical functionalities.

### 2.3. Hyperdimensional Computing Model

This section details the core components of the HDC framework as applied in this study, including the architectural principles behind the HDC model and the strategies for encoding tokens into high-dimensional vectors.

#### 2.3.1. Design Principles of a Vector Symbolic Architecture

HDC, also known as Vector-Symbolic Architectures (VSA), is an emerging computational paradigm that leverages high-dimensional vectors to represent and process information [16,18]. Unlike traditional computing which relies on precise, low-dimensional data, HDC operates on vectors with thousands of dimensions, where individual elements carry little meaning but the overall pattern and relationships between vectors are highly significant. Importantly, HDC can successfully represent non-numerical, categorical, and temporal information thanks to its symbolic representation capability. HDC typically employs either binary hypervectors (logic-1 and logic-0, particularly valuable for hardware implementations) or bipolar hypervectors (+1 and −1, commonly used in software platforms) as its atomic datatype. The core principles that enable HDC to perform cognitive tasks, such as learning and classification, are:

- <u>High dimensionality</u>: information is encoded into vectors of very high dimensionality (e.g., 1,000 dimensions). Such long vectors can holistically represent multiple inputs, even when they come from different data formats, by embedding them into a single unified datastructure. This high dimensionality provides a vast space for representing complex data and allows for robust and fault-tolerant operations;
- <u>Distributed representation</u>: concepts and data are represented as distributed patterns across the entire hypervector. This means that information is not localized to a single bit or small group of bits, making the representation resilient to noise and errors;
- <u>Associative memory</u>: HDC inherently supports *associative memory*, where similar inputs map to similar hypervectors. This property is crucial for tasks like classification, as it allows for the comparison of new, unseen data (from *item memory*) with learned patterns;
- <u>Vector operations</u>: HDC relies on a set of fundamental algebraic operations that manipulate these high-dimensional vectors. These operations, such as *binding, bundling*, and *permutation*, allow for the encoding of relationships, the aggregation of information, and the creation of new, meaningful representations, and together are known as the MAP (Multiply-Add-Permute) model:
  ∘ <u>Binding:</u> A key operation that combines two hypervectors into a merged representation. Binding can be performed using element-wise multiplication for bipolar vectors (+1/–1) or logic XOR for binary vectors. This operation produces a new hypervector that encodes the association of the two input vectors. Importantly, the original hypervectors can be approximately recovered from the bound result if one of the components is known.
  ∘ <u>Bundling:</u> An operation (i.e., element-wise addition in bipolar space or population count in binary space) that aggregates multiple hypervectors into a single representative hypervector. Bundling is typically used to summarize information across features or samples. For example, in image processing, hypervectors of pixel intensity values after being bound with their corresponding positional hypervectors are bundled across all pixels to form a single hypervector representation. Bundling across samples of the same class yields a prototype hypervector that summarizes the overall class.
  ∘ <u>Permutation:</u> A scrambling operation (i.e., circular shift) that reorders the elements of a hypervector to encode order or temporally relevant information of each hypervector. For instance, in language processing, atomic symbols (letters) represented as hypervectors can be permuted according to their position within a word, thereby preserving the sequential structure. Permutation also helps generate nearly orthogonal hypervectors, which improves independence among tuples and strengthens encoding capacity.

These design principles collectively enable HDC architectures to learn from data, make classifications, and exhibit properties akin to human memory and cognition, all while maintaining a high degree of robustness and efficiency.

#### 2.3.2. Hypervector Encoding Strategy

Because SMILES structures are inherently symbolic, HDC provides a naturally compatible framework for encoding them. After the tokenization of a SMILES string, the resultant tokens from each of the three strategies previously presented are transformed into high-dimensional bipolar vectors. This fundamental HDC encoding translates symbolic information into a numerical format suitable for high-dimensional yet simple arithmetic for a single pass learning method. The specific encoding approach varied depending on the tokenization strategy to preserve or abstract positional information as required. For all tokenization strategies, we employed the random projection encoding method: each unique token identified across the entire dataset for a given tokenization strategy is assigned to a *unique, randomly generated*, and *high-dimensional* bipolar vector. These base hypervectors are sparse and balanced, containing equal numbers of +1 and −1 entries to maximize orthogonality. In probabilistic terms, the likelihood of a position taking a +1 or −1 value is approximately 0.5, representing an unbiased distribution between the minimum (*P* = 0) and maximum (*P* = 1) probabilities. The dimensionality of the generated hypervectors is typically set to *D* = 10, 000, which provides sufficient representational capacity and robustness.

In case of an atom-wise and k-mer-based tokenization, in order to preserve the sequential order of tokens within a SMILES string, a positional encoding scheme is applied. After generating a base hypervector for each unique token, the hypervector for a token at a specific position within a SMILES sequence is permuted. Specifically, the hypervector of the token at position *i* (where the first token is at position 0) is circularly shifted by *i* positions. This permutation operation effectively binds the token’s identity with its positional context. Once all tokens in a SMILES string are encoded with their respective positional permutations, the hypervectors are all bundled together into a single, composite hypervector representing the entire chemical compound. For instance, if a SMILES string is tokenized into a sequence of tokens *T*_0_, *T*_1_, *T*_2_, …, *T*_*N*_ with *N* number of tokens, and their corresponding base hypervectors are 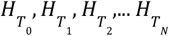, then the compound hypervector *H*_*C*_ is derived a 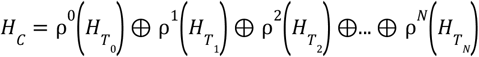, where 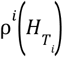 denotes a circular shift of the hypervector *H* by *i* positions, and ⊕ represents the bundling operation.

On the other hand, in case of a fragment-based tokenization, where the order of fragments is implicitly captured by the fragment definitions themselves, the hypervector for an entire SMILES string is constructed by directly bundling the base hypervectors of its constituent fragments. The overall encoding procedure is outlined in Algorithm 1.

##### Algorithm 1:. Pseudocode for the compound hypervector encoding process. The function takes a SMILES string and an encoding type as input and returns a single composite hypervector (HC) by bundling the permuted (for atom-wise and k-mer-based tokenizations) or base (for fragment-based tokenization) hypervectors of its constituent tokens

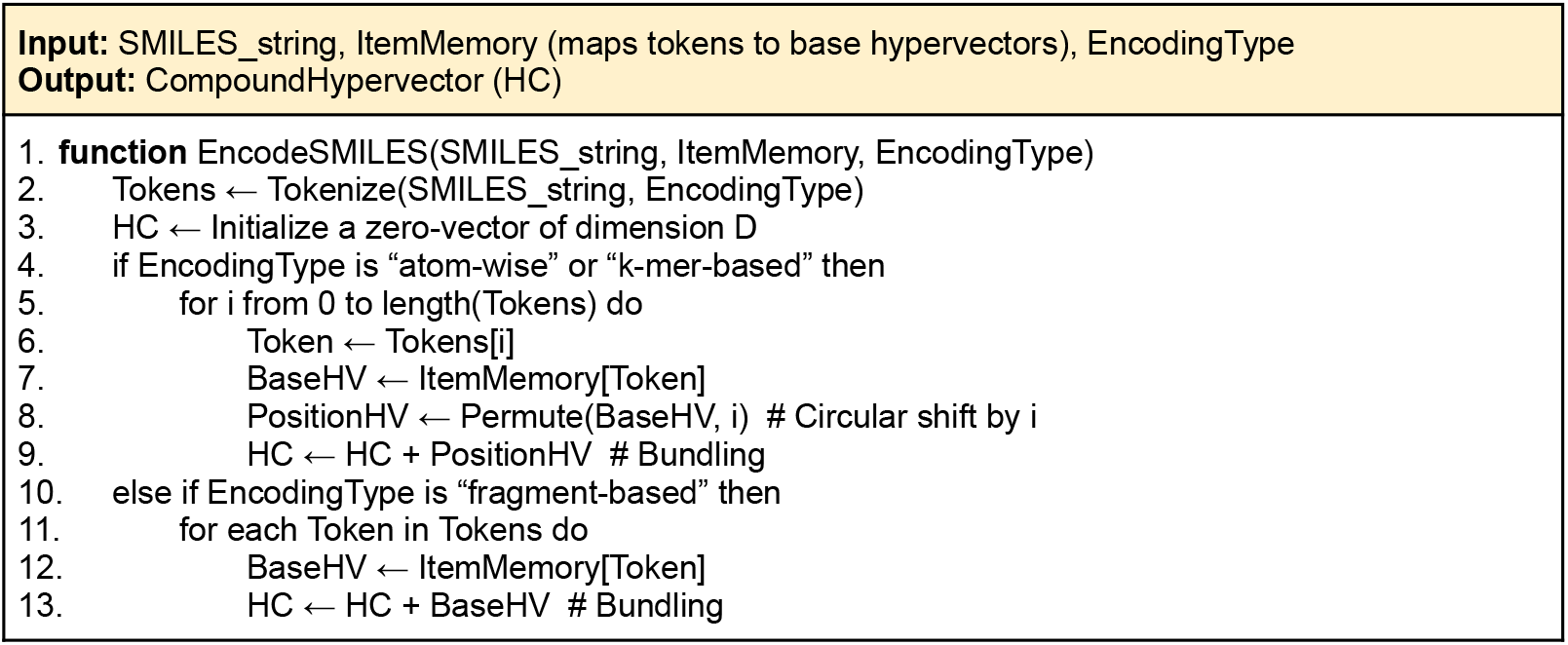
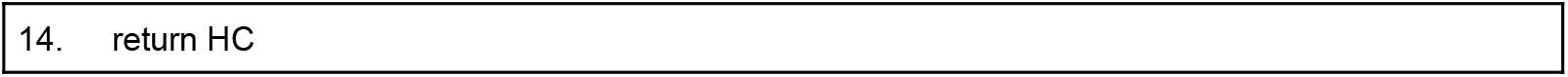

#### 2.3.3. Classification Model Components

The HDC model implemented for predicting the toxicity of chemical compounds consists of several key components that facilitate the learning and inference processes:

- <u>Item memory</u>: this serves as a dictionary or lookup table that stores the unique base hypervectors for all individual tokens (atoms, k-mers, or fragments) identified during the tokenization phase. Each unique token is mapped to a distinct, randomly generated bipolar hypervector of a chosen dimensionality *D*. This memory is static once initialized and provides the fundamental building blocks for constructing compound hypervectors;
- <u>Associative memory (class prototype)</u>: for each of the 12 receptors, an associative memory is constructed. This memory stores a prototype hypervector for each class (toxic and non-toxic). These class prototypes are learned during the training phase and represent the aggregated characteristics of all compounds belonging to a specific class for a given receptor. The prototype for a class is formed by bundling all the individual compound hypervectors that belong to that class in the training set;
- <u>Encoder</u>: the encoder module is responsible for taking a raw SMILES string and, based on the chosen tokenization strategy (atom-wise, k-mer, or fragment-based), generating its corresponding compound hypervector. This involves tokenizing the SMILES string, retrieving or permuting the base hypervectors from the item memory, and then building them according to the rules defined above;
- <u>Classifier</u>: this module performs the inference step. For an unseen compound’s hypervector, it calculates its cosine similarity to each of the class prototypes (toxic and non-toxic) stored in the associative memory for the specific receptor being evaluated. The compound is then classified into the class whose prototype hypervector exhibits the highest similarity to the compound’s hypervector.

#### 2.3.4. Error Mitigation Process

While HDC models are inherently robust and can achieve good performance with a single pass through the training data (a process that takes the name of one-shot learning), their accuracy can often be further refined through iterative training. This iterative process serves as a crucial error mitigation mechanism, particularly beneficial when dealing with noisy or imbalanced datasets.

During each training iteration, the class prototypes within the associative memory are updated. This update mechanism involves presenting training samples to the model and adjusting the respective class prototype hypervectors based on the classification outcome. Specifically, if a training sample is correctly classified, its hypervector reinforces its assigned class prototype. Conversely, if a training sample is misclassified, its hypervector can be used to slightly adjust the prototypes to minimize future misclassifications. This error mitigation process encompasses two key concepts of HDC’s usage: one is incremental learning, which reinforces the correctly labeled class dynamically during training. The second approach is to dynamically unlearn the misclassified training sample from the trained model by simply applying a reverse operation, i.e., subtraction. Compared to traditional learning systems, this approach is cost-effective for both incremental and unlearning concepts, utilizing algebraic operations only. The proposed iterative refinement helps to:

- <u>Reduce noise sensitivity</u>: by repeatedly exposing the model to training data, the prototypes become more robust to individual noisy samples, as the collective influence of correctly classified samples gradually outweighs the impact of outliers;
- <u>Improve class separation</u>: over a few iterations (epochs), the prototypes for different classes (toxic and non-toxic) become more distinct and representative of their respective categories, leading to clearer boundaries in the high-dimensional space. This enhances the model’s ability to differentiate between classes during inference;
- <u>Address data imbalance</u>: in datasets where one class is significantly more prevalent than another, initial prototypes might be biased towards the majority class. Iterative updates can help the model to better learn the characteristics of the minority class.

The number of training iterations is empirically determined to achieve optimal performance and converge to a minimal error rate, close or ideally equal to 0. The retraining procedure is detailed in Algorithm 2.

##### Algorithm 2:. Pseudocode for the iterative model retraining procedure. The function initializes class prototypes using one-shot learning and then refines them over multiple epochs by adjusting the prototypes based on misclassifications to minimize the training error rate

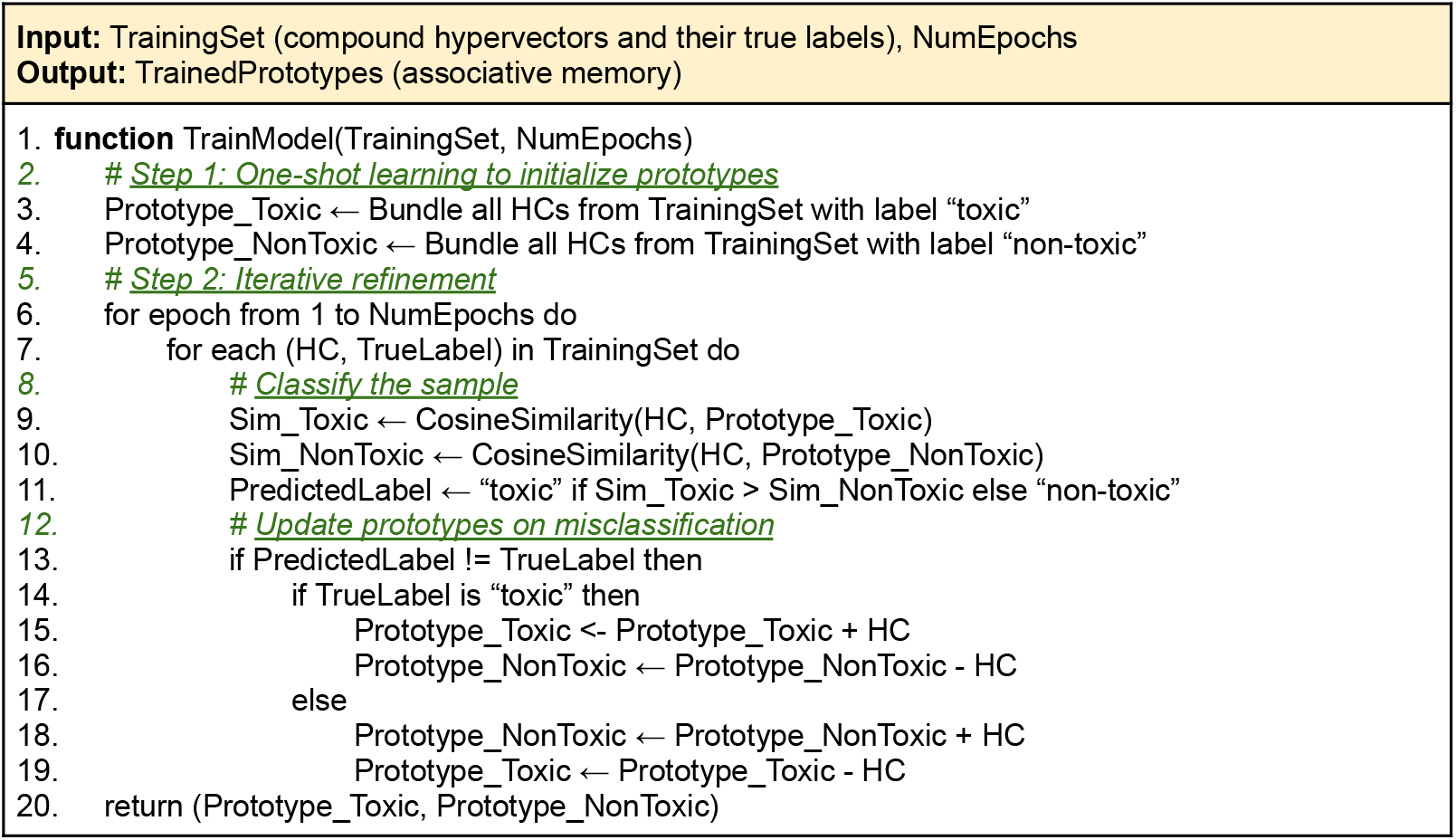

To provide a holistic view of the entire experimental workflow, from model creation to the classification of new compounds, the complete process is summarized in Figure 1. This flowchart illustrates the distinct training and prediction phases, integrating the concepts detailed in Algorithm 1 and 2 into a single, cohesive architecture.

**Figure 1:**
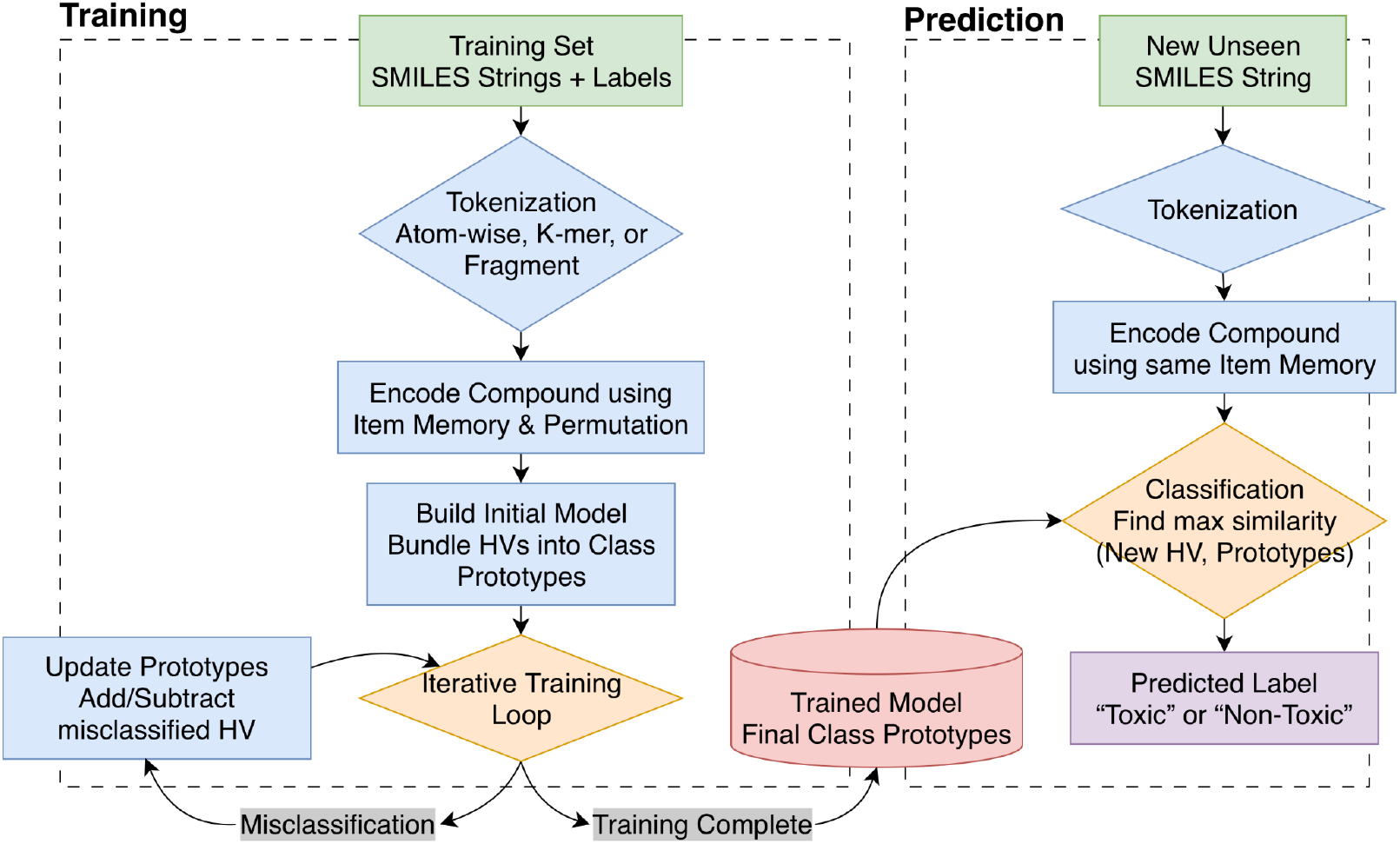
Flowchart of the complete VSA-based toxicity prediction pipeline. The diagram is divided into two main phases: (i) the <u>Training</u> phase begins with the labeled training dataset, proceeds through tokenization and encoding (as detailed in Algorithm 1), and uses an iterative refinement loop (as detailed in Algorithm 2) to produce the final, trained class prototypes; (ii) the <u>Prediction</u> phase shows how a new, unlabeled SMILES string is processed through the same encoding pipeline and then classified by comparing its resulting hypervector against the trained prototypes to determine the most likely class.

## 3. RESULTS

This section presents the empirical findings from our evaluation of the HDC model for toxicity prediction. The computational pipeline was constructed primarily using the *hdlib* library v0.1.21 [27] for the core HDC implementation. We first detail the experimental setup, our strategy for handling class imbalance, and the evaluation metrics. We then analyze the impact of k-mer length on model performance to identify an optimal configuration. Subsequently, we provide a comprehensive comparison analysis of the three primary tokenization strategies (i.e., atom-wise, k-mer-based, and fragment-based) across all 12 toxicity targets in the Tox21 dataset. This tokenization was performed using *SmilesPE* v0.0.3 [28] for the atom-wise and k-mer-based approaches, and the *RDKit* library [29] release 2025.03.5 to extract chemically meaningful fragments for the fragment-based approach. We then demonstrate the effectiveness of the iterative error mitigation process, analyze the computational performance, and provide a contextual comparison against traditional, descriptor-based machine learning methods.

### 3.1. Experimental Setup and Evaluation Metrics

All experiments were conducted using hypervectors of dimensionality *D* = 10, 000, a dimension shown to provide sufficient capacity for robust information encoding. The entire pipeline was developed in Python 3, and the computational tasks were executed on a workstation equipped with 4 Intel Xeon Platinum 8276 L Central Processing units (CPUs) (112 cores/224 threads) @ 2.20GHz and 6TB of Random Access Memory (RAM) running CentOS Linux 7. The dataset for each of the 12 biological receptors was partitioned into a training set (80%) and test set (20%).

A significant challenge in the Tox21 dataset is the severe class imbalance for individual receptor targets, where the number of non-toxic compounds vastly exceeds the number of toxic compounds as shown in Table 1. To address this problem, we implemented a repeated undersampling and averaging strategy. For a given receptor, let *X* be the number of compounds in the minority (toxic) class and *Z* be the number in the majority (non-toxic) class. We calculated the ratio *N* = *Z*/*X* and randomly partitioned the *Z* non-toxic compounds into *N* distinct, non-overlapping subsets. Subsequently, we performed *N* independent training and evaluation runs in 5-folds cross-validation. In each run, the full set of *X* toxic compounds was combined with one of the *N* subsets of non-toxic compounds to create a nearly perfectly balanced training dataset.

A complete HDC model was trained on each of these *N* balanced datasets, and its performance was evaluated on the test set. The final performance metrics reported for that receptor (i.e., Accuracy, Precision, Recall, and F1-score) are the average of the scores obtained from all *N* models. This entire procedure was repeated independently for all 12 receptors. This approach ensures that the model is evaluated against all non-toxic samples while mitigating the bias introduced by class imbalance during training. Given this setup, the F1-score is emphasized as the primary indicator of robust performance. The entire process of building, training, and evaluating the models for all 12 receptors, including all undersampling runs in 5-folds cross-validation, considering all 3 tokenization methods for a total of 2,960 classification models, was completed in approximately 2 hours and 23 minutes, underscoring the computational efficiency of the overall methodology.

### 3.2. Optimization of k-mer length

To determine the optimal configuration of the k-mer-based tokenization strategy, we systematically evaluated model performance using different k-mer lengths, specifically for *k* values ranging from 2 to 10. The objective was to identify a length that effectively captures meaningful local structural motifs from the SMILES strings without becoming overly specific or sparse.

Our analysis revealed a clear and consistent trend: shorter k-mer lengths yielded superior predictive performances. As illustrated in Figure 2, the highest average F1-score across all toxicity targets was achieved with *k* = 4, reaching a value of *∼*64%.

**Figure 2:**
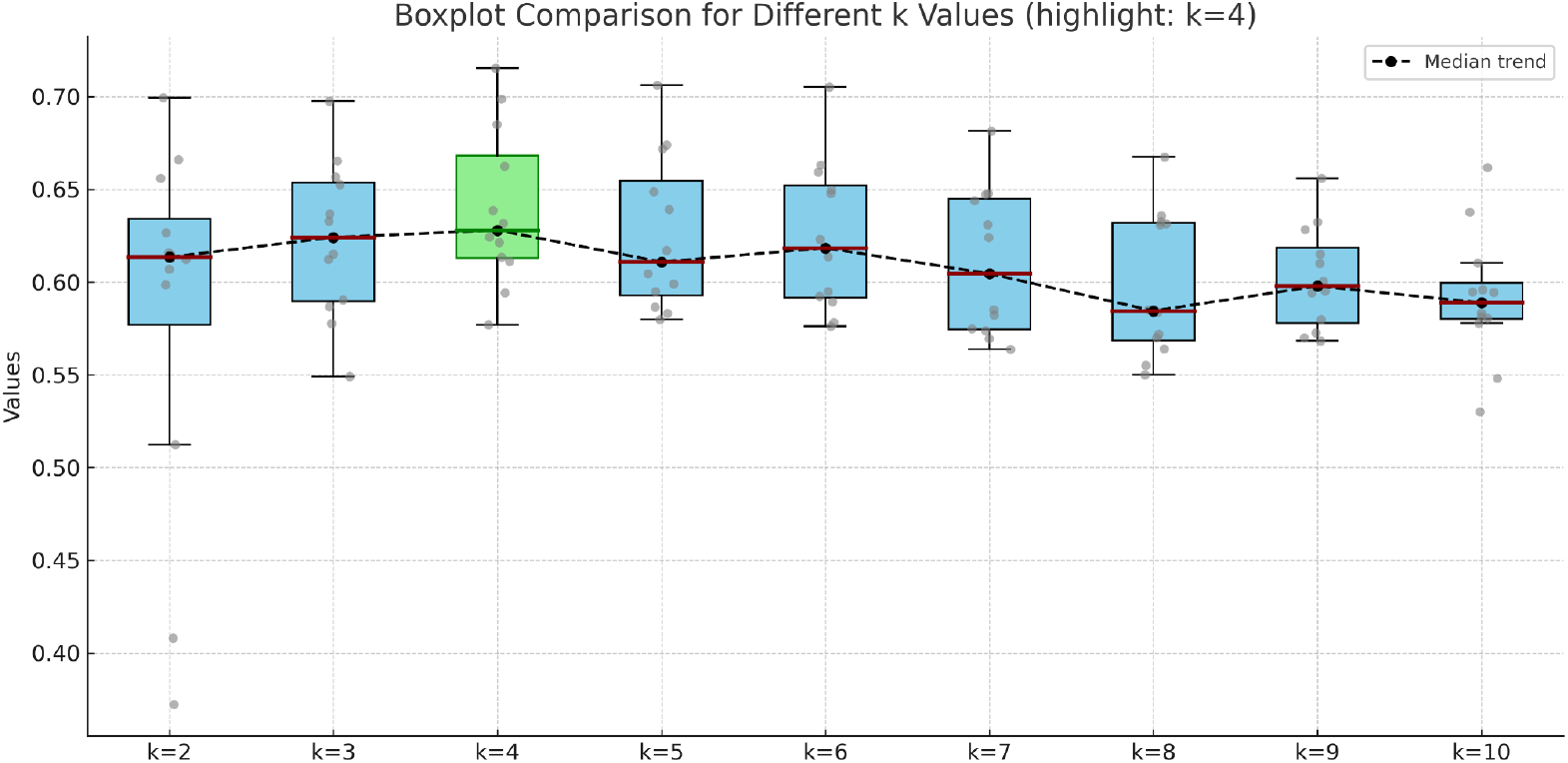
Boxplot built over the F1-scores computed on all 12 HDC models (one model per receptor) considering *Y* different partitions, each of them evaluated in 5-folds cross-validation, highlighting *k* = 4 as the top performing one (colored in green).

As the value of *k* is increased, a gradual but steady decline in performance was observed. For instance, with *k* = 5, the average F1-score dropped to 62%, and for *k* = 8 it further decreased to 59%. This performance degradation suggests that while longer k-mers capture more extended local contexts, they do so at the cost of generality. Longer sequences are statistically rarer, leading to a sparser vocabulary of tokens and reducing the chance of finding recurring, predictive patterns between the training and testing sets. In contrast, a k-mer length of 4 appears to strike an effective balance, creating tokens that are complex enough to represent local chemical environments (e.g., *C*(= *O*)) but common enough to be generalizable. Based on these findings, a k-mer length of *k* = 4 was selected as the optimal value and used for all subsequent comparative analyses.

### 3.3. Comparative Analysis of Tokenization Strategies

The main aim of our study was to compare the efficacy of different structural representation granularities for HDC-based toxicity prediction. We evaluated the performance of models built using *atom-wise, fragment-based*, and the *optimized k-mer-based* (*k* = 4) tokenization strategies. Table 2 summarizes the average classification performance across all 12 Tox21 targets for each method.

**Table 2:**
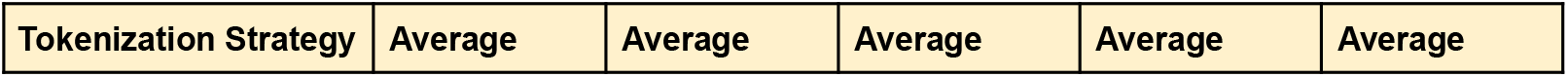
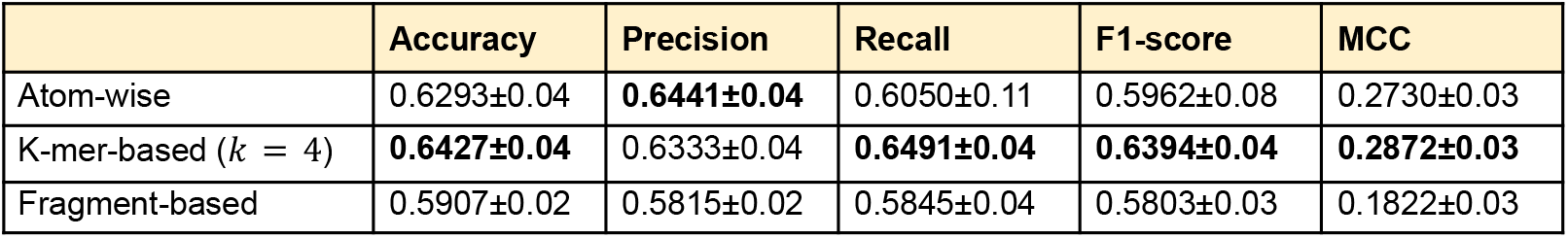
Average classification performance of different tokenization strategies reporting accuracy, precision, recall, F1-score, and Matthews Correlation Coefficient (MCC) with their standard deviations.

| Tokenization Strategy | Average | Average | Average | Average | Average |
| --- | --- | --- | --- | --- | --- |

|  | Accuracy | Precision | Recall | F1-score | MCC |
| --- | --- | --- | --- | --- | --- |
| Atom-wise | 0.6293±0.04 | **0.6441±0.04** | 0.6050±0.11 | 0.5962±0.08 | 0.2730±0.03 |
| K-mer-based ($k = 4$) | **0.6427±0.04** | 0.6333±0.04 | **0.6491±0.04** | **0.6394±0.04** | **0.2872±0.03** |
| Fragment-based | 0.5907±0.02 | 0.5815±0.02 | 0.5845±0.04 | 0.5803±0.03 | 0.1822±0.03 |

The complete, disaggregated performance data for each of the *N* undersampling runs, from which the averages in Table 2 were derived, are provided for atom-wise, k-mer-based, and fragment-based strategies in Supplementary Spreadsheets S1, S2, and S3, respectively.

The aggregated results clearly indicate that the k-mer-based (*k* = 4) tokenization strategy slightly outperformed the other two methods, achieving the highest average F1-score of *∼*64%. This performance suggests that capturing local, overlapping character sequences within the SMILES string provides an effective balance of granularity and contextual information for the HDC model.

In contrast, the atom-wise and fragment-based tokenization methods yielded the lowest performance, with an average F1-score of *∼*59% and *∼*58% respectively. The performance of the atom-wise models is significant as it validates the hypothesis that the most granular representation of a chemical structure does not contain sufficient information for an effective classification. On the other hand, the performance of the fragment-based models is equally significant because, while this approach is designed to capture chemically meaningful functional groups, its effectiveness appears limited by the predefined nature of its fragment library. This method is inherently biased towards known chemistry and may fail to represent novel structural motifs or subtle atomic configurations that are critical for determining toxicity which the other, more agnostic methods successfully capture.

To provide a more nuanced view, Table 3 presents the F1-scores for all the 12 toxicity targets, showcasing how performance can vary depending on the biological endpoint.

**Table 3:**
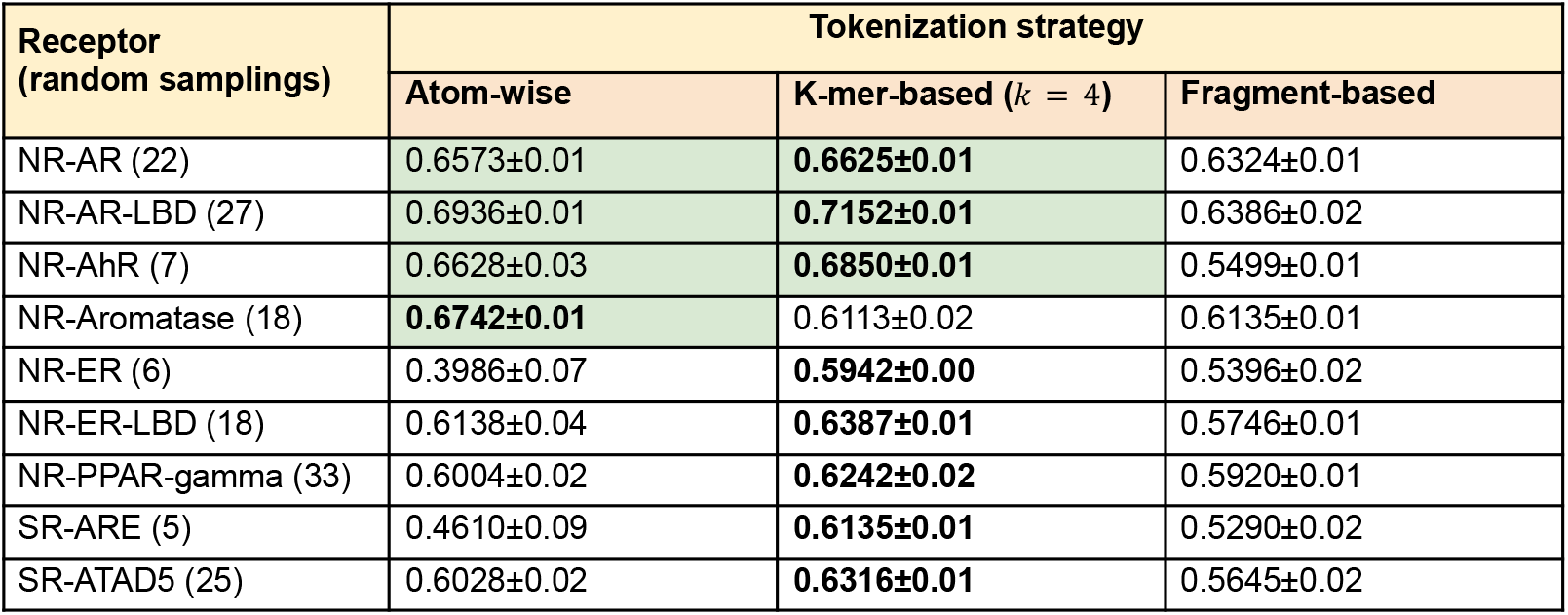
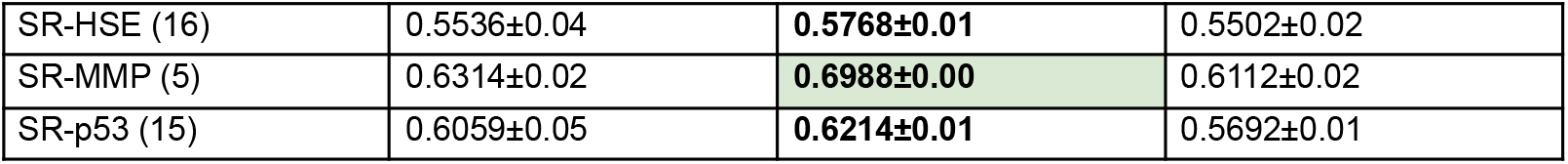
Comparison of F1-scores for the three tokenization strategies on the 12 Tox21 receptors. Each value is derived from a two-level averaging process: for each of *N* random sampling runs, a 5-fold cross-validation was performed. The reported score is the final average of the F1-scores from all *N* runs. The number of runs *N* varies for each receptor and it is defined as the number of non-toxic compounds divided by the number of toxic compounds. Scores exceeding 65% are highlighted in green.

For the NR-AR-LBD and SR-MMP receptors, the k-mer-based approach showed a distinct advantage, suggesting that specific local substructures, rather than just individual atoms or large functional groups, are key drivers of activity. Interestingly, for the NR-AR-LBD receptor, all models performed relatively well, with the fragment-based model being more competitive compared to the other models built with the same tokenization method for the other receptors. This may indicate that toxicity for this specific receptor is strongly linked to the presence of a few well-defined, reactive functional groups that are adequately captured by the fragment library.

### 3.4. Effectiveness of the Iterative Error Mitigation Process

To evaluate the efficacy of the iterative training procedure as an error mitigation mechanism, we monitored the model’s training error rate at each iteration up to a maximum number of 100 iterations. The *training error rate* is defined as the sum of false positives and false negatives divided by the total number of samples in the training set. This metric provides a direct measure of how well the model’s class prototypes have learned to represent the training data.

The results showed a rapid and consistent reduction in training error across all tokenization strategies and toxicity targets. Considering all models built in this study, we reached an initial error rate of *∼*40% in the worst case. As the iterative process began, the class prototypes were progressively refined, leading to a steep decline in the error rate. For the majority of models, the training error rate converged to a value at or near 0%. This convergence indicates that the iterative updates successfully adjusted the prototype hypervectors to correctly classify nearly all samples in the training set. This process effectively strengthens the representation of each class, improving the separation between them in the high-dimensional space and making the final model more robust. The findings confirm that iterative training is a crucial step for optimizing the HDC model and minimizing classification errors learned from the data.

### 3.5. Analysis of Computational Performance

A key motivation for exploring HDC is its potential for computational efficiency. We measured the total training time for a single balanced model and the average inference time per compound for each tokenization strategy.

The k-mer-based model (*k* = 4) was the fastest to train, taking 1.85 seconds. The atom-wise and fragment-based models required slightly more time, taking 8.81 and 5.10 seconds respectively.

However, all individual models were trained in under 10 seconds, a time drastically lower than that required for training other conventional machine learning models. Most importantly, the inference time for all models was negligible, taking a few milliseconds per compound.

To further characterize the computational footprint, we also monitored the resource utilization for a single model run. The primary memory consumer was the *item memory*, which stores the token-to-hypervector mappings. Consecutively, the peak RAM usage was directly proportional to the vocabulary size of the tokenization strategy. The k-mer-based (*k* = 4) model, with its large vocabulary of unique 4-mers, required the most memory, typically utilizing around 750MB of RAM. The fragment-based and atom-wise models, with their more constrained vocabularies, were significantly leaner, requiring approximately 300MB and less than 100MB of RAM, respectively. In all cases, the training process for a single model was computationally lightweight, utilizing a single CPU core.

This demonstrates the profound scalability of the HDC approach, making it an exceptionally well-suited component for high-throughput screening protocols of massive chemical libraries.

### 3.6. Contextual Comparison with Descriptor-Based Methods

To contextualize our findings, it is important to compare our approach with conventional machine learning models that represent the state-of-the-art in QSAR. Therefore, we compare our descriptor-free HDC approach with conventional machine learning models. The comparative analysis was streamlined using PyCaret [30], an automated machine learning framework that facilitates the rapid training and evaluation of multiple models. These methods, such as Random Forest, Support Vector Machines, or Deep Learning, typically rely on a pipeline where pre-computed molecular descriptors serve as input features. These descriptors (i.e., numerical features), are engineered to explicitly quantify the molecule’s topological, physiochemical, and electronic properties. They are designed to capture specific characteristics believed to govern biological activity [31–33]. A comprehensive feature set in our study includes 219 descriptors, categorized as molecular properties (physicochemical), electronic/charge descriptors, topological indices, geometric/shape descriptors, fingerprint-based density features, constitutional descriptors (counts), functional group counts (fragments) and structural/ring system descriptors, were calculated based on the *RDKit* library’s built-in functionality.

Once this feature matrix is constructed, it serves as the input for training the machine learning models. As has been extensively reported in literature [14,34–37] and is confirmed by our benchmark analysis shown in Figure 3, such descriptor-based models can achieve very high predictive performance on the Tox21 dataset, with F1-scores often ranging from 59% to 84%.

**Figure 3:**
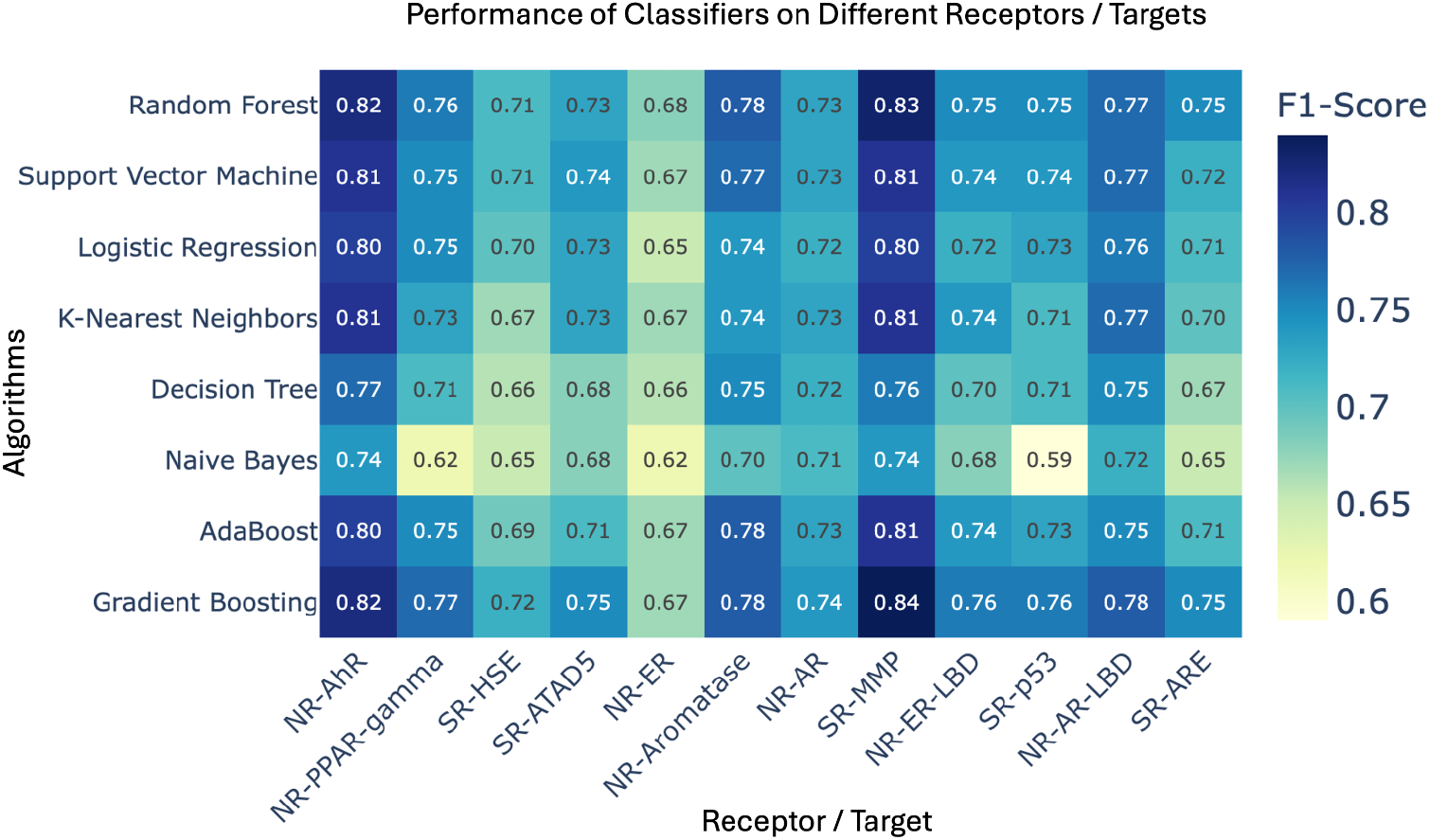
Comparative heatmap of F1-scores for state-of-the-art QSAR-based models trained on molecular descriptors. This figure illustrates the predictive performance of several conventional machine learning algorithms on the 12 individual toxicity prediction tasks from the Tox21 dataset. Each cell in the heatmap represents the F1-score achieved by a specific algorithm on a specific receptor target, with all results derived from a 5-fold cross-validation. Unlike the HDC approach, these models were trained on a feature set of pre-calculated chemical descriptors.

While these scores are often higher than the 71.52% F1-score achieved by our best-performing k-mer-based HDC model, this comparison highlights a fundamental trade-off in methodology. The often superior performance of conventional models is contingent upon an extensive and computationally intense feature engineering step, the calculation of hundreds or thousands of chemical descriptors. This process requires significant domain expertise and separates the structural representation from the learning algorithm. This extensive feature engineering step, however, introduces significant complexity and a substantial computational cost. On average, the training and 5-fold cross-validation time for these models ranged from a best case of 3.47 minutes for Naive Bayes to a worst case of 22.73 minutes for AdaBoost.

In contrast, our HDC-based approach achieves a competitive F1-score of 71.52% through a much simpler, end-to-end pipeline that operates directly on the raw SMILES strings, with an average training time of less than 15 seconds. The value of our method, therefore, lies not in surpassing the raw accuracy of highly optimized, descriptor-based systems, but in demonstrating a fundamentally different paradigm that offers an exceptional balance of good predictive power, radical simplicity, and profound computational efficiency. This makes the HDC approach particularly attractive for scenarios requiring rapid, large-scale screening where the overhead of feature engineering is prohibitive. When it comes to screening millions of virtual molecules, including enumerated libraries and previously extracted patterns, HDC can be an alternative for making a first-pass toxicity prediction, discarding obvious candidates in milliseconds before the descriptor-heavy QSAR validation. Since descriptor-dependent calculation is slow and expensive, the proposed approach provides a computationally efficient alternative.

## 4. DISCUSSION AND CONCLUSIONS

Here, we interpret the empirical findings presented in the previous section, contextualizing them within the broader field of computational toxicology. We discuss the significance of our results, from the proof-of-concepts for the descriptor-free paradigm to the practical implications of the trade-off between computational efficiency and predictive accuracy, before concluding with the limitations of this study and future research.

### 4.1. A proof-of-concept for a descriptor-free paradigm

The results of this study serve as a critical proof-of-concept in uniting HDC with the challenging domain of chemical toxicity prediction. Our findings establish a viable, descriptor-free pipeline and, while the absolute predictive performance does not surpass that of highly optimized, state-of-the-art methods, the work provides valuable insights into the methodology and its trade-offs. The primary contribution of this research is not the creation of a new top-performing model, but rather the systematic exploration of a novel computational paradigm with *efficiency* considerations, and the clear trends that emerged from it.

### 4.2. The importance of local chemical context in Hyperdimensional Computing encoding

The central finding of our work is the performance of the k-mer-based tokenization strategy, particularly with an optimal k-mer length of 4. This result is highly informative, suggesting that the most predictive structural information for toxicity is contained not merely in individual atoms or in large, predefined functional groups, but in the local chemical context and short-range (nearest neighbor) atomic arrangements. The k-mer approach, by creating overlapping sequences of characters, effectively captures these local motifs (e.g., *C*(= *O*)*N*, representing part of an amide group) without being constrained by a fixed chemical dictionary. It strikes an optimal balance between the high granularity of the atom-wise method and the high-level abstraction of the fragment-based method. This balance allows the model to learn a rich vocabulary of structural patterns that are directly relevant to biological interactions, supporting our second conclusion that the choice of tokenization strategy is critical for achieving optimal performance.

The performance of the atom-wise and fragment-based methods, while lower, provides essential context. Their results serve as a valuable baseline, demonstrating that simply encoding individual atoms or pre-defined fragments is less effective. This reinforces the importance of the local context captured by k-mers and highlights a potential limitation of using a fixed chemical dictionary (fragments) which may lack the flexibility to represent all relevant toxicophores. This comparative analysis leads to our second conclusion: the choice of tokenization strategy is a critical determinant of model performance in HDC-based cheminformatics.

### 4.3. The trade-off: radical simplicity and efficiency vs. accuracy

One of the most significant outcomes of this study is the stark contrast in computational philosophy. Conventional QSAR models achieve higher accuracy, but this performance is contingent upon a complex, multi-stage pipeline involving the calculation of hundreds of chemical descriptors. Our work highlights a fundamental trade-off. This leads to our third conclusion: the HDC paradigm offers a compelling balance of moderate predictive power, radical pipeline simplicity, and profound computational efficiency. The ability to train models in seconds and perform inference in milliseconds directly from raw SMILES strings makes this approach exceptionally suitable for rapid, large-scale preliminary screening, where the cost and complexity of feature engineering would be prohibitive.

### 4.4. Limitations and future directions

It is crucial, however, to acknowledge the limitations of the study. The primary limitation is the moderate predictive accuracy of the model when compared to descriptor-based systems. This indicates that while our descriptor-free approach captures essential information, it does not yet encapsulate the full spectrum of physicochemical properties that determine biological activity. Nevertheless, it opens a new and fruitful avenue in cheminformatics by emphasizing computation-efficient design. Secondly, while the HDC encoding process is transparent, interpreting the final, aggregated class prototype hypervectors to extract human-understandable chemical rules remains a significant challenge. Despite these limitations, this work, being the first of its kind, poses an important future research pathway for the integration of HDC with toxicity prediction. Future research should focus on bridging the performance gap. This could involve developing more sophisticated HDC encoding schemes that might incorporate physicochemical properties without full descriptor calculation, or exploring hybrid models that combine the speed of HDC for an initial screening with more complex models for subsequent refinement. The lightweight and highly parallel nature of HDC operations makes this especially attractive for application-specific and portable device design. In scenarios where millions of virtual molecules must be screened (e.g., drug discovery), or where on-site chemical safety monitoring requires immediate triage, HDC offers a computationally efficient alternative capable of discarding toxic candidates in milliseconds.

### 4.5. Conclusions

In conclusion, this work successfully validates HDC as a promising and computationally efficient paradigm for toxicity prediction. Future research should focus on bridging the performance gap. This could involve developing more sophisticated HDC encoding schemes that might incorporate physicochemical properties without full descriptor calculation, or exploring hybrid models that combine the speed of HDC for an initial screening with more complex models for subsequent refinement. Such advancements will be critical in developing the next generation of tools for enhancing our ability to ensure human and environmental health.

## ADDITIONAL INFORMATION

### Availability

The *hdlib* Python library (v0.1.21) [27] used for building the VSA in this study is an open-source tool and is freely available under the MIT license. The source code is available at https://github.com/cumbof/hdlib and it can be installed via the Python Package Index (PyPI – *pip install hdlib*) and Conda (*conda install -c conda-forge hdlib*). *SmilesPE* (v0.0.3) and *RDKit* (release 2025.03.5) are also open-source and their source code is available on GitHub at https://github.com/XinhaoLi74/SmilesPE and https://github.com/rdkit/rdkit and can be installed via the Python Package Index (*pip install SmilesPE rdkit*) and Conda as well on the Bioconda [38] channel (*conda install -c bioconda SmilesPE rdkit*).

The specific pipeline developed for the classification of chemical compounds based on their SMILES representations, along with the preprocessed Tox21 dataset used in this research, are also publicly available under the examples directory of the *hdlib* GitHub repository at https://github.com/cumbof/hdlib/tree/main/examples/tox21.

## Abbreviations

CPU: Central Processing Unit
HDC: Hyperdimensional Computing
MAP: Multiply-Add-Permute
NR-AR: Nuclear Receptor - Androgen Receptor
NR-AR-LBD: Nuclear Receptor - Androgen Receptor Ligand Binding Domain
NR-AhR: Nuclear Receptor - Aryl Hydrocarbon Receptor
NR-Aromatase: Nuclear Receptor - Aromatase
NR-ER: Nuclear Receptor - Estrogen Receptor alpha
NR-ER-LBD: Nuclear Receptor - Estrogen Receptor Ligand Binding Domain
NR-PPAR-gamma: Nuclear Receptor - Peroxisome Proliferator-Activated
PyP: Python Package Index
QSAR: Quantitative Structure-Activity Relationship
RAM: Random Access Memory
SMILE: Simplified Molecular Input Line Entry System
SR-ARE: Stress Response - Antioxidant Response Element
SR-ATAD5: Stress Response - ATPase Family AAA Domain
SR-HSE: Stress Response - Heat Shock Factor Response
SR-MMP: Stress Response - Mitochondrial Membrane Potential
SR-p53: Stress Response - p53 Pathway
Tox21: Toxicology in the 21st Century
VSA: Vector-Symbolic Architecture

## Author Contribution

FC and JJ conceived the research; FC and SA designed the classification model and implemented the encoding and classification pipeline; FC, KD, and JJ performed the analysis; JJ, BR, DC, and DB analyzed the results and provided insights from a biochemical perspective; DB supervised the research; FC, KD, JJ, BR, DC, SA, and DB wrote the manuscript and agreed with its final version.

## Acknowledgments

We would like to acknowledge the use of AI in refining the clarity and readability of this manuscript. The AI assistance was primarily used for tasks such as sentence restructuring, word choice suggestions, and identifying potentially unclear phrasing. We emphasize that the AI was used solely for language enhancement and did not contribute to the generation of research ideas, data analysis, or the interpretation of results. All conclusions drawn and insights presented in this manuscript are solely the product of the authors’ own analysis and expertise.

